# Mechanism of heme binding by CP motifs in the BACH1 DNA-binding region

**DOI:** 10.64898/2026.08.28.747782

**Authors:** Yubin Huang, Louise Fairall, Fred Muskett, Cyril Dominguez, Andrew Hudson, John W.R. Schwabe

## Abstract

BACH1 is a heme-regulated “basic-leucine-zipper” containing transcriptional repressor that binds its DNA recognition elements as a heterodimer with MAFK. Heme-binding is thought to be mediated by several cysteine-proline (CP) motifs and to trigger dissociation of the heterodimer from DNA. However, the structural basis of heme recognition and the mechanism of heme-binding and heme-mediated DNA dissociation remains unresolved. We have used UV-visible spectroscopy, 2D-NMR and DNA-binding assays to explore both heme-binding and DNA dissociation of a minimal BACH1 construct containing 2 CP motifs (C492(CP5) and C646(CP6)) flanking the DNA-binding domain. We find that heme is able to bind to both CP motifs, but also to other non-CP cysteines and histidines in the construct. Using NMR spectroscopy, we identify a structured binding pocket in which heme interacts with both C646(CP6) and Cys621. However, DNA-binding assays show that C646(CP6) is not required for heme-mediated DNA dissociation of the BACH1:MAFK heterodimer. Using UV-visible spectroscopy we show that C492(CP5) also recruits heme with a second ligand, a conserved histidine, H559, in the BACH1 DNA-recognition helix. Mutation of C492(CP5) reduces but does not abolish heme-mediated dissociation from DNA. Together, our findings reveal the structural basis for heme recognition by BACH1 and the mechanism, for heme-mediated dissociation from DNA.

## Introduction

BTB and CNC homology 1 (BACH1) is a heme binding transcriptional repressor belonging to the Cap ‘n’ Collar basic-leucine-zipper (CNC-bZIP) family (1, 2). It plays roles in multiple physiological processes including heme metabolism, oxidative stress response, cell cycle, inflammation, mitochondrial metabolism and macrophage adaptation (3–8).

BACH1 is a cysteine-rich 736 amino acid protein with two structured domains – an amino-terminal BTB domain and a carboxy-terminal bZIP DNA-binding domain (Figure 1A). The BTB domain of BACH1 mediates homo-dimerisation (9, 10) and is thought to recruit HDAC1 (histone deacetylase 1) as part of the NuRD complex leading to transcriptional repression (8, 11). The bZIP domain mediates heterodimerisation with the small musculoaponeurotic fibrosarcoma (MAF) proteins including MAFF, MAFG and MAFK (12, 13). The BACH1:MAFK heterodimer binds to multiple Maf recognition elements (MAREs) in oxidative stress-response genes resulting in repression of genes including heme oxygenase-1 (HO-1) and NADPH quinone oxidoreductase 1 (NQO1) (14–16).

**Figure 1.**
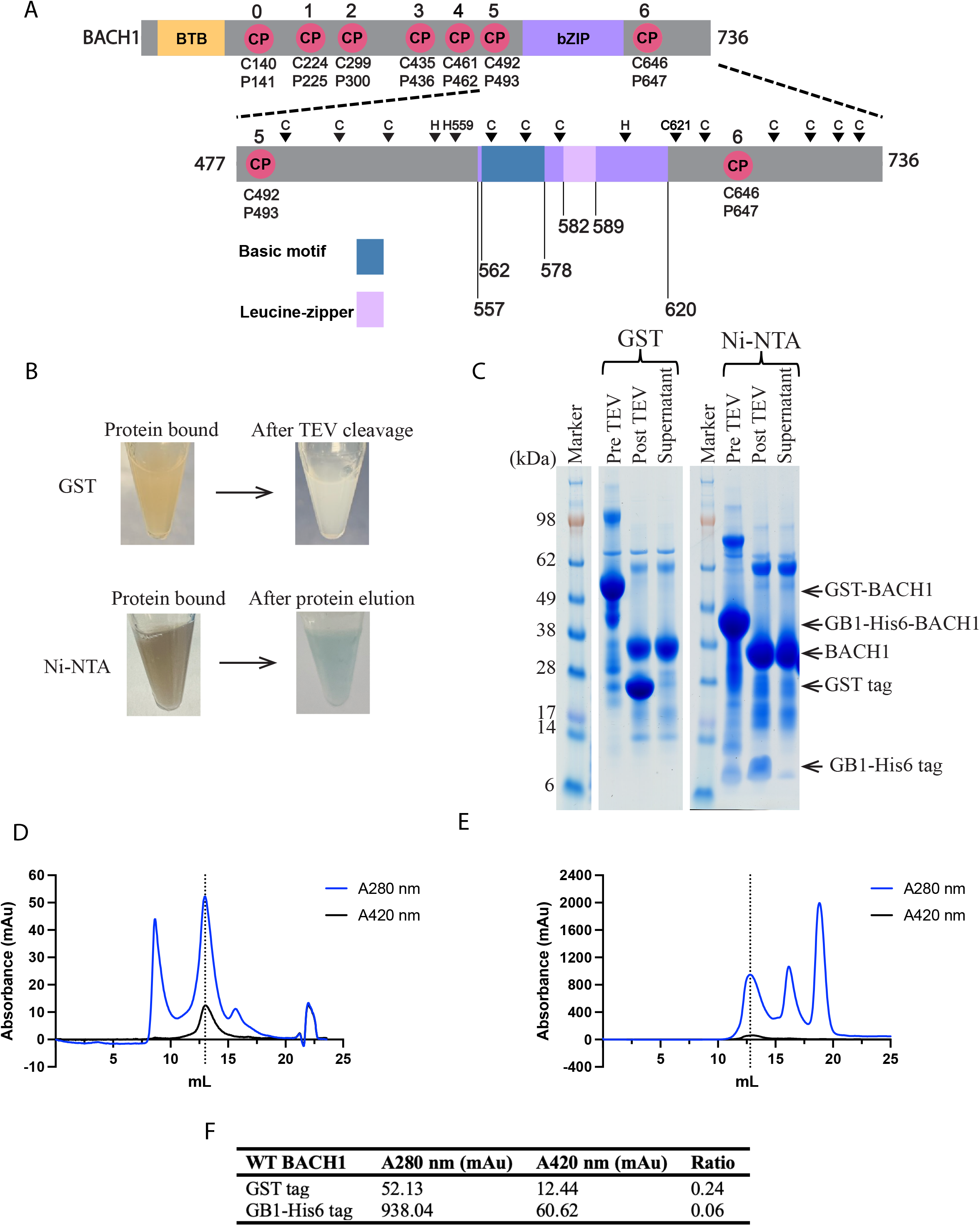
BACH1 co-purifies with endogenous heme. **(A)** Schematic of human full-length BACH1 showing BTB domain, bZIP DNA binding domain and 7 CP motifs, and BACH1 (aa: 477-736) with two CP motifs, other 12 non-CP cysteines and 3 histidines indicated. (**B)** Brown colouration of BACH1 (aa: 477-736) bound to Glutathione Sepharose resin and BACH1 bound to Ni-NTA resin. The resins are restored to the original colour after TEV cleavage. **(C)** SDS-PAGE analysis of GST and Ni-NTA affinity purification. Pre TEV (affinity tagged BACH1), Post TEV (after TEV cleavage) and supernatant samples were loaded on the gel. **(D)** Size exclusion chromatogram of cleaved GST-tagged BACH1 and (**E)** cleaved GB1-His6-BACH1. A280 nm (blue) and A420 nm (black). BACH1 protein peak is indicated by a dashed line. **(F)** The A_420_/A_280_ ratios of cleaved GST-tagged and cleaved GB1-His6 tagged BACH1.

Under conditions of elevated heme, such as those arising from oxidative stress or tissue damage, free heme binds directly to BACH1 in the nucleus and causes the dissociation of the repressive BACH1:MAFK heterodimer from DNA (2). A different heterodimer of MAFK with Nuclear factor erythroid 2-related factor 2 (NRF2) – i.e. MAFK:NRF2 – binds to MARE sequences in place of BACH1:MAFK resulting in activation of genes such as HO-1 expression (4, 17, 18). This mutually exclusive binding of BACH1:MAFK and MAFK:NRF2 provides reciprocal regulation of the oxidative stress-response.

In addition to causing dissociation of the BACH1:MAFK heterodimer from DNA, heme-binding to BACH1 is also thought to result in nuclear export and thereby promote interaction of the BTB domain with the E3 ligases (SCF^FBXO22^ and SCF^FBXL17^) resulting in BACH1 degradation (19, 20).

Human BACH1 contains 34 cysteine residues with 7 arranged as cysteine–proline (CP) dipeptide motifs (Figure 1A). By analogy with the CP heme-recognition motifs in proteins such as Heme Activator Protein 1 (HAP1), it is widely believed that the CP motifs mediate interaction with heme (21, 22). CP3 and CP4 motifs have been shown to be important for heme-induced nuclear export by Exportin-1 (23). It has been suggested that heme binding to C492(CP5) and C646(CP6) motifs, which flank the DNA-binding bZIP domain, might induce conformational changes, leading to DNA dissociation and heterodimer destabilisation (24, 25). Recently, it has also been observed that heme binding to C649 (CP6) directly recruits another E3 ligase (CRL2^FEM1B^) for BACH1 degradation (26). This is in addition to the BTB domain E3 ligase interactions.

To understand the mechanism through which heme binds to BACH1 and modulates DNA-binding activity we expressed a construct containing the bZIP domain and the two flanking CP motifs C492(CP5) and C646(CP6). A combination of difference UV-VIS spectroscopy and 2D-NMR approaches have been used to investigate heme-binding to wild-type (WT) and mutant BACH1, which has led us to identify two distinct heme binding sites. The first is a pre-formed carboxy-terminal pocket that coordinates heme via C646(CP6) and C621. In contrast, the second is a conformationally-inducible amino-terminal pocket that forms upon heme binding, where the heme is coordinated by C492(CP5) and H559. Electrophoretic

Mobility Shift assays (EMSA) show that, as expected, WT BACH1:MAFK dissociates from DNA at heme concentrations >20 µM. A BACH1:MAFK heterodimer in which all histidines and cysteines in BACH1 are mutated to asparagine and serine respectively remains associated with DNA at heme concentrations > 100 µM. A BACH1:MAFK heterodimer lacking C646(CP6) dissociates from DNA exactly like WT suggesting that the C646(CP6):C621 pocket is not required for heme mediated DNA-dissociation. In contrast, a heterodimer lacking C492(CP5) shows impaired, but not abolished heme-mediated DNA-dissociation, suggesting that heme binding to other non-CP ligands may also contributes to heme-mediated dissociation.

## Results

### BACH1 co-purifies with endogenous bacterial heme

To investigate how heme interacts with BACH1, and how heme modulates BACH1-DNA binding, we expressed a minimal BACH1 construct (aa 477-736) that contains the bZIP DNA-binding domain flanked by two CP motifs (C492(CP5) & C646(CP6)). We chose this construct since previous work suggested that C492(CP5) & C646(CP6) might be sufficient to regulate DNA binding activity (2, 24). In addition to these CP motifs, this construct of BACH1 contains three histidines (His) and twelve non-CP cysteines (Cys), any of which could also coordinate to heme (Figure 1A).

We initially expressed a GST-tagged BACH1 (aa: 477-736). Upon purification, the protein exhibited a brown colouration on glutathione Sepharose resin (Figure 1B). Following removal of the GST tag and further purification using size-exclusion chromatography the protein retained a Soret absorption peak (B band) at 420 nm (Figure 1C & D). This suggests that the construct acquired endogenous heme during the bacterial expression. A GB1-His6-tagged version of the equivalent construct of BACH1 also exhibited a Soret band following tag removal and purification. However, the relative intensity of this peak compared to the protein was much lower (Figure 1B, C & E), likely because the very much higher expression level exceeded the bacterial heme pool. Based on the A_420_/A_280_ ratios (Figure 1F), we estimate that the cleaved GB1-His6-BACH1 represents a “low-heme” sample with >75% of the protein lacking bound heme.

### Spectroscopic discrimination of heme coordination by histidine vs cysteine

To probe heme coordination in more detail, we used difference UV-visible spectroscopy with the “low-heme” BACH1 (aa 477-736). Experiments were performed with equimolar protein and heme (200 µM each). Under these conditions, multiple Lewis-basic residues (His and Cys) within the construct will compete for heme binding, with the equilibrium favouring occupation of the highest-affinity site by a single heme molecule on each protein molecule in the equimolar mixture. Heme may bind either a single His or Cys residue (yielding a 5-coordinate complex) or two residues (yielding a 6-coordinate complex). All measurements were carried out at pH 8.0 in the presence of 5mM dithiothreitol (DTT) to both mimic the reducing environment of the nucleus and avoid the formation of di-sulphide bonds. Under these mildly basic conditions, roughly one-third of cysteine thiol side-chains are expected to be deprotonated. Once coordinated to a heme iron, a deprotonated thiol is likely to be fully stabilised. DTT can itself act as an axial ligand to heme, via one of its thiolate groups, and therefore can occupy any open axial site on a 5-coordinate protein-heme complex as has been observed in crystal structures of heme oxygenase and cytochrome P450 (PDB: 3I9T & 6IAO) (Supplementary Figure S1) (27, 28).

Difference spectra were generated by subtracting the spectrum of heme alone (with DTT) from a heme:BACH1 complex. The observed difference spectrum reflects a convolution of the spectra of heme coordinated by different combinations of Cys, His and DTT. The difference spectrum for WT BACH1 (aa 477-736) showed two peaks in the Soret region of approximately equal intensity at around 370 nm and 420 nm (Figure 2A). Note that a weak Q-band at 550 nm and a charge-transfer (CT) band at 650 nm are also observed (29). To confirm that these features arise specifically from heme interaction with Cys or His residues, we generated a variant in which all Cys and His residues were replaced with serines and asparagines, respectively. The difference spectrum lacked any distinct Soret peak, and showed only a broad feature around 400nm, consistent with the absence of defined heme coordination (Figure 2B).

**Figure 2.**
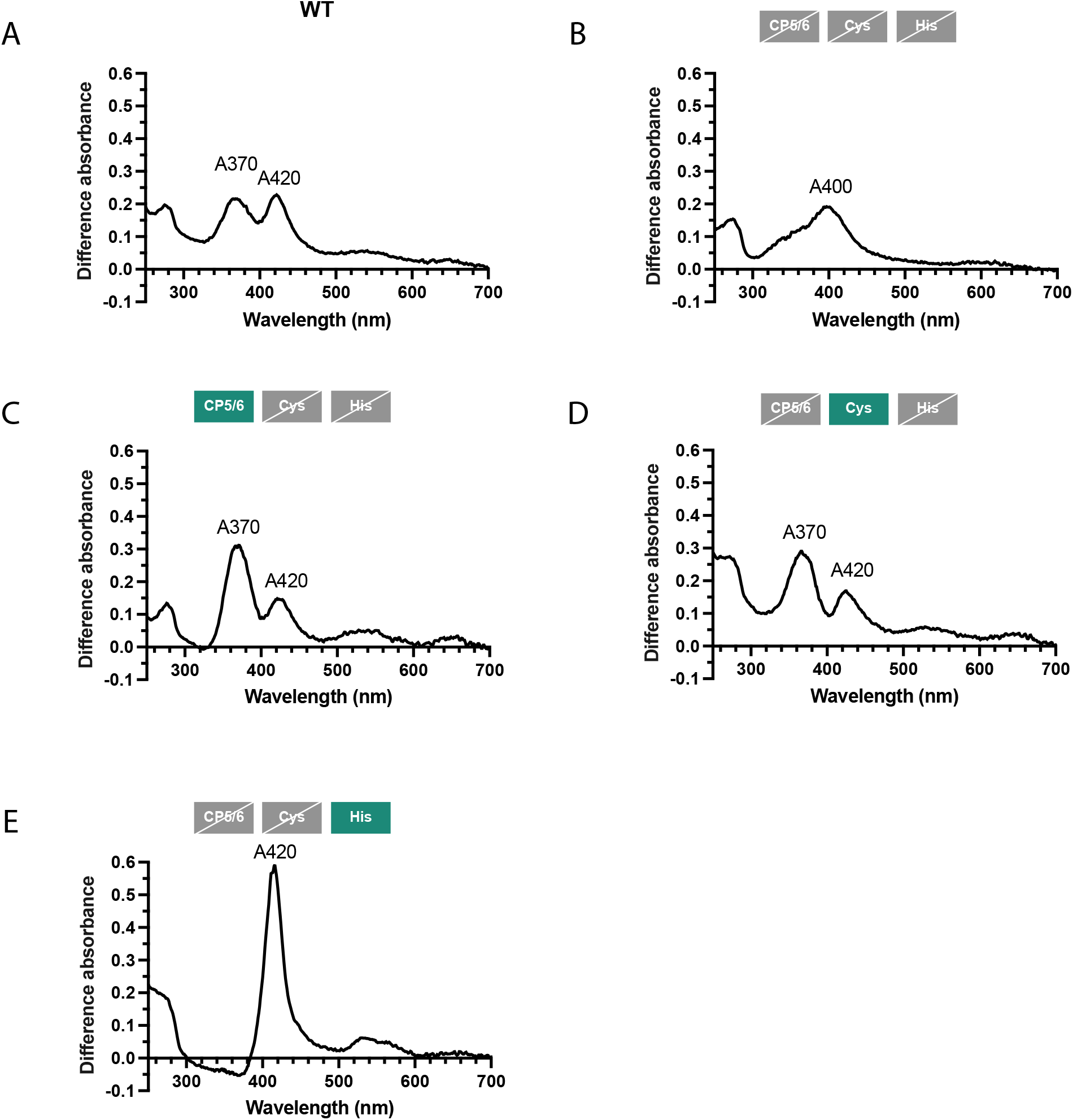
Difference UV-visible spectra of heme bound BACH1. (A-E) Spectra of WT BACH1 and mutant BACH1. Three boxes on the top of each spectrum indicate the status of two CP cysteines, non-CP cysteines and histidines residues: green boxes denote retained residues, while grey boxes crossed by a slash denote mutated residues.

To determine whether C492(CP5) and C646(CP6) are specific contributors to heme binding, we prepared a construct in which all other cysteines and histidines were mutated. The resulting difference spectrum exhibited peaks at 370 nm and 420 nm in an approximate 2:1 ratio (Figure 2C). It would appear that this 2:1 ratio is likely to be diagnostic of heme binding to Cys residues with, in this case, both C492(CP5) and C646(CP6) contributing a single thiolate ligand and with the opposite axial site likely filled by a DTT thiolate. The simultaneous appearance of peaks at around 370 and 420 nm is the hallmark of a hyperporphyrin spectrum, which arises when a heme is bound strongly to two thiolate ligands (in this case, one from Cys and another from DTT) in a six-coordinate geometry. The 370 nm feature corresponds to a thiolate-metal charge-transfer band mixed into the Soret manifold, whereas the 420 nm feature is the porphyrin B band.

To compare heme binding between CP motif and other cysteines, we next generated a construct in which all the CP cysteines and all histidines were mutated, leaving only the non-CP cysteines intact. This construct produced a difference spectrum essentially identical to that of the CP-only mutant, including the 370nm:420nm ratio of approximately 2:1 (Figure 2D). This suggests that spectroscopically, all cysteines in the BACH1 construct behave similarly with respect to heme coordination and that the hyperporphyrin signature is consistent with a Cys/DTT axial pair.

In contrast, a BACH1 construct retaining only the three His residues, with all Cys residues mutated to serines, has a very distinct difference spectrum comprising a single strong peak in the Soret region at 420nm (Figure 2E). This is characteristic of histidine coordination. In a mono-His complex, the second axial site would again be occupied by DTT, but DTT-thiolate coordination is too weak to induce a hyperporphyrin spectrum alone, consistent with the absence of a 370 nm feature. Note that the absence of a CT band, is also characteristic of histidine coordination.

Taken togther, these results allow us to distinguish the coordination modes through binding to Cys (2:1 370nm:420nm) or His residues (strong 420nm only).

### A structured heme-binding pocket involving C646(CP6) and C621

To further explore heme-binding to the various BACH1 constructs we collected (^1^H-^15^N)-TROSY spectra for a number of BACH1 constructs. The spectrum of a truncated WT *apo*-BACH1 construct comprising residues 477-736 contained 100 resolved peaks compared with 246 expected resonances arising from non-proline residues (Supplementary Figure S2). The ^1^H chemical shift dispersion is rather limited (7.5 – 8.6 ppm) as might be expected for an intrinsically disordered / helical protein. The failure to observe 146 peaks is likely to be due to peak overlap and or exchange broadening. 12 peaks were observed with an ^15^N chemical shift between 108-112 ppm and ^1^H shift between 8.0-8.5 ppm. These peaks are likely to correspond to the 12 glycine residues in this truncated BACH1 construct.

(^1^H-^15^N)-TROSY spectra were also collected for two other truncated *apo*-BACH1 constructs (with residues 477-613 & 548-736) (Supplementary Figure S3). Although these truncated proteins showed some aggregation, in contrast to the longer protein, it was possible to assign many peaks to the amino-terminal region, carboxy-terminal region or the common central region (Supplementary Figure S4).

Interestingly, the addition of heme to WT BACH1 (477–736), in a 1:1 ratio, resulted in some very specific changes in the NMR spectrum: at least 15 peaks disappeared (broadened or shifted) and 13 new peaks were observed for *holo*-BACH1. Of the 15 peaks that are clearly broadened or shifted, 13 are from the amino-terminal sequence of 70 amino acids (Figure 3A).

**Figure 3.**
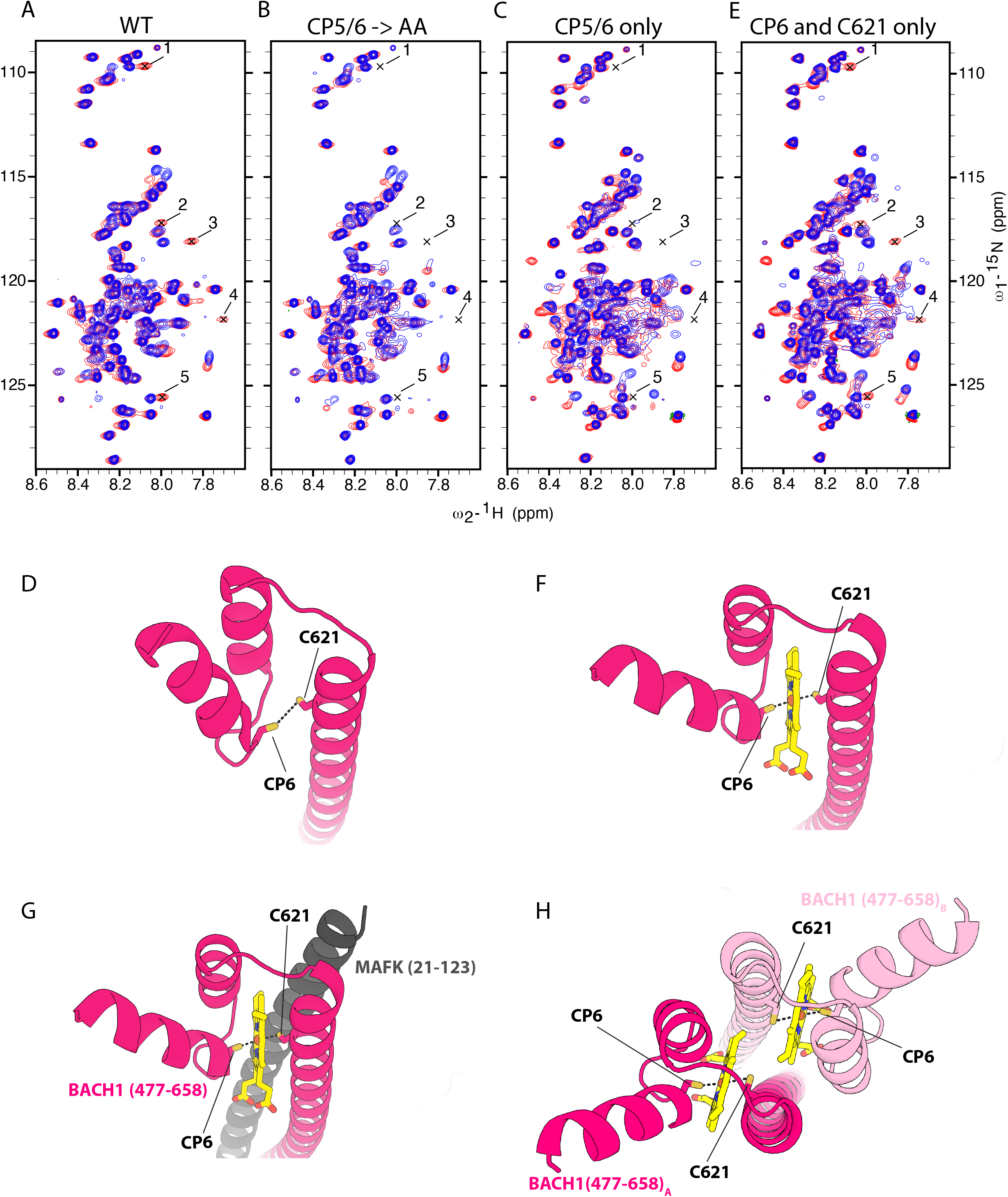
C646(CP6) and C621 are involved in a specific heme binding pocket. (^1^H-^15^N)-TROSY spectra overlay of BACH1 with (red) and without (blue) heme: **(A)** WT BACH1, **(B)** BACH1 lacking two CP motifs (each CP mutated to AA), **(C)** BACH1 mutant containing only CP5 and CP6 and **(E)** BACH1 mutant containing only CP6 and C621. Five CP-related peak changes on heme-binding are indicated by number 1-5. **(D)** AlphaFold 3 model of apo BACH1 (aa: 477-658). (**F)** 6-coordinated heme formed by C646(CP6) and C621 from BACH1 monomer. **(G)** C646(CP6) and C621 can form 6-coordination with heme in BACH1:MAFK heterodimer. **(H)** Two hemes engaging CP6 and C621 as axial ligands in each monomer of a BACH1:BACH1 homodimer.

To determine whether these spectral changes are the result of the CP motifs interacting with the heme, we prepared a BACH1 construct lacking C492(CP5) and C646(CP6) (each CP mutated to AA). There were rather few changes in the spectra of *apo*-form of these mutant proteins compared with WT *apo*-BACH1 – only the loss of two of the peaks from the WT spectrum and the appearance of two new peaks (Supplementary Figure S5). However, when heme was added to the (C492(CP5) & C646(CP6) → AA) mutant protein, we observed fewer changes than seen with the WT protein allowing us to define five specific, CP-related, changes on heme-binding that are observed in the WT protein but lost following mutation of the C492(CP5) & C646(CP6) motifs (see 1-5 in Figures 3A & B).

To confirm that these changes are due to the interaction of C492(CP5) & C646(CP6) interaction with heme, we created a mutant protein containing only C492(CP5) and C646(CP6) - with all other cysteine and histidine residues mutated to serine and asparagine respectively. To our surprise none of the five CP-related changes were restored suggesting that the changes may be dependent not only on the CP motifs but also on other non-CP heme ligands (Figure 3C).

An AlphaFold-predicted structure of BACH1 suggests that C621 is positioned to potentially act as a 2nd protein ligand to make a 6-coordinate complex for the iron when heme is bound to the C646(CP6) motif (Figure 3D) (30). To test this hypothesis, we created a mutant BACH1 protein containing only C646(CP6) and C621. All other cysteines were mutated to serine and all three histidines to asparagine. On adding heme to this protein, all five CP-related changes were restored (Figure 3E).

In conclusion, these data provide strong evidence that there is a specific and pre-formed heme-binding pocket involving C646(CP6) and C621. Binding to this pocket accounts for the majority of the specific changes in the (^1^H-^15^N)-TROSY spectra. Modelling heme into this binding pocket using AlphaFold 3 and Phenix(31), reveals a non-polar cavity in which heme can be readily accommodated between the two thiolate ligands from C646(CP6) and Cys621 (Figure 3F). Importantly, this pocket can accommodate heme in both a BACH1:MAFK heterodimer and a BACH1 homodimer (Figures 3G & H).

### A second heme-binding site involving H559 and C492(CP5)

To investigate whether C492(CP5) might also contribute to heme recognition, as has been previously suggested, we created a different series of BACH1 mutants. Compared with WT protein, a mutant lacking C492(CP5) showed increased absorbance at 370nm i.e. a shift toward cysteine, and away from histidine, interactions (Figure 4A & B). This was unexpected, given that we had removed a cysteine residue, and implies that C492(CP5) must be further assisting heme coordination to a histidine residue. In contrast, a construct lacking C646(CP6), but including CP5, showed a dramatically-increased peak at 420nm suggesting an increase in histidine interactions with heme (Figure 4B) – this was not unexpected given that C646(CP6) supports the formation of a bis-cysteine complex with heme.

**Figure 4.**
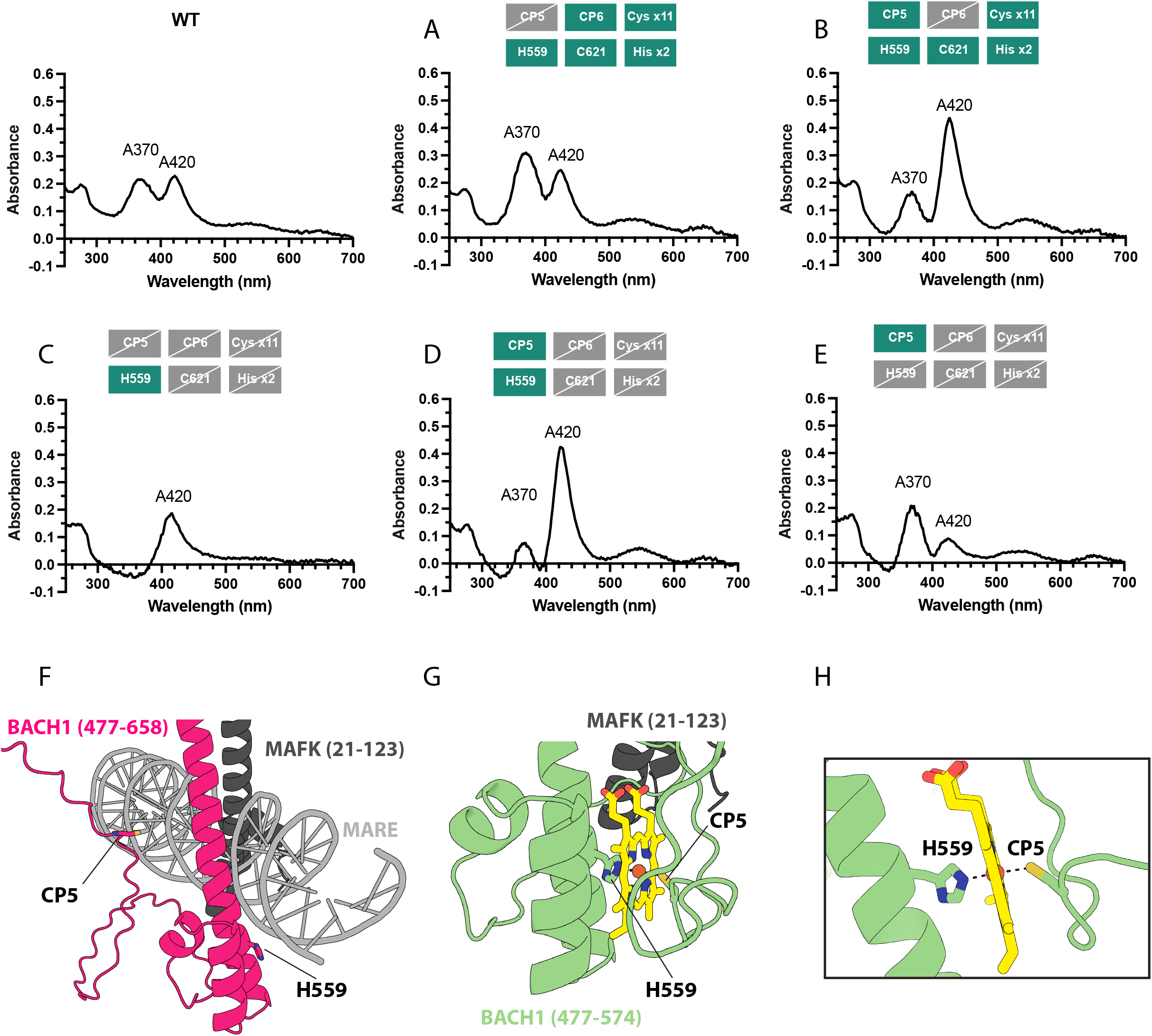
CP5 and H559 form another heme-binding pocket. (A-E) Difference UV-visible spectra of heme bound BACH1 mutants. Six boxes on the top of each spectrum indicate the status of two CP motifs, H559, C621, other non-CP eleven cysteines and other two histidines residues: green boxes denote retained residues, while grey boxes crossed by a slash denote mutated residues. All spectra were measured with addition of heme to the BACH1 (at 1:1 ratio) in 20 mM Tris-HCl pH 8.0 buffer with 50 mM NaCl and 5 mM DTT. **(F)** AlphaFold 3 model of apo BACH1:MAFK heterodimer with MARE. **(G)** AlphaFold 3 model of heme bound BACH1 (C492(CP5) and H559) (aa: 477-574) with MAFK (aa: 21-123). **(H)** Zoomed-in view of (G).

We speculated that H559, located in the DNA-binding helix, might compete strongly for heme binding. However, a construct containing just H559 showed a modest peak at 420nm indicating the likelihood of only a weak heme interaction (Figure 4C). Remarkably, a construct containing H559 together with C492(CP5) showed a greatly increased 420nm peak suggesting that C492(CP5) strongly supports heme interaction with H559 (Figure 4D). As would be expected, a control construct containing only C492(CP5) shows cysteine-only interaction (approximately 2:1 ration for A370:A420; Figure 4E).

Homologous to other bZIP proteins, H559 is located in the DNA-binding helix of BACH1, oriented toward the DNA and likely making a contact with the phosphate backbone in the DNA-bound complex. In contrast to H559, AlphaFold 3 suggests that C492(CP5) is located in a disordered region (Figure 4F).

AlphaFold prediction of the amino-terminal region of the BACH1 construct (477–574), in the presence of a heme ligand, suggests that the disordered region may reposition so as to form a conformationally-induced heme-binding site, in which H559 and C492(CP5) form a 6-coordinate heme complex (Figure 4G & H). Given that H559 is close to the basic region that mediates DNA-interaction (Figure 4F), we hypothesised that heme-binding at this site might explain how heme is able to dissociate the BACH1:MAFK heterodimer from DNA. Interestingly, we note that the disordered region also contains additional cysteine residues (C496, C515 & C522). It is possible that these could serve as alternative heme ligands, together with H559, to form 6-coordinate heme complexes and have the same impact on heterodimer dissociation from DNA.

### Role of heme binding in dissociation of the BACH1:MAFK complex from DNA

To explore whether heme-binding to the BACH1 (aa:477-736) protein mediates dissociation of a BACH1:MAFK heterodimer from DNA, we expressed and purified a BACH1(aa:477-736):MAFK(aa:21-123) heterodimer (Supplementary Figure S6) and used electromobility - shift assays to observe DNA-binding to a MARE. Having prepared stoichiometric complexes of wild-type and mutant proteins with the MARE (at 0.5 µM), we added increasing amounts of heme. The wild-type complex dissociates around 20 µM heme (pseudo IC50 = 23 µM) (Figure 5A & E). In contrast, a mutant heterodimer, in which all histidine and cysteine residues within BACH1 were mutated to asparagine and serine, remained bound to the MARE at > 100 µM heme (Figure 5B). Interestingly, a complex in which BACH1 C646(CP6) was mutated dissociated with a pseudo IC50 of 26 µM (Figure 5C & E) suggesting that C646(CP6) does not play an important role in heme-mediated dissociation from DNA. In contrast, a BACH1:MAFK heterodimer in which the BACH1 C492(CP5) motif was mutated to AA shows impaired dissociation from DNA, requiring a heme concentration > 57 µM (Figure 5D & E).

**Figure 5.**
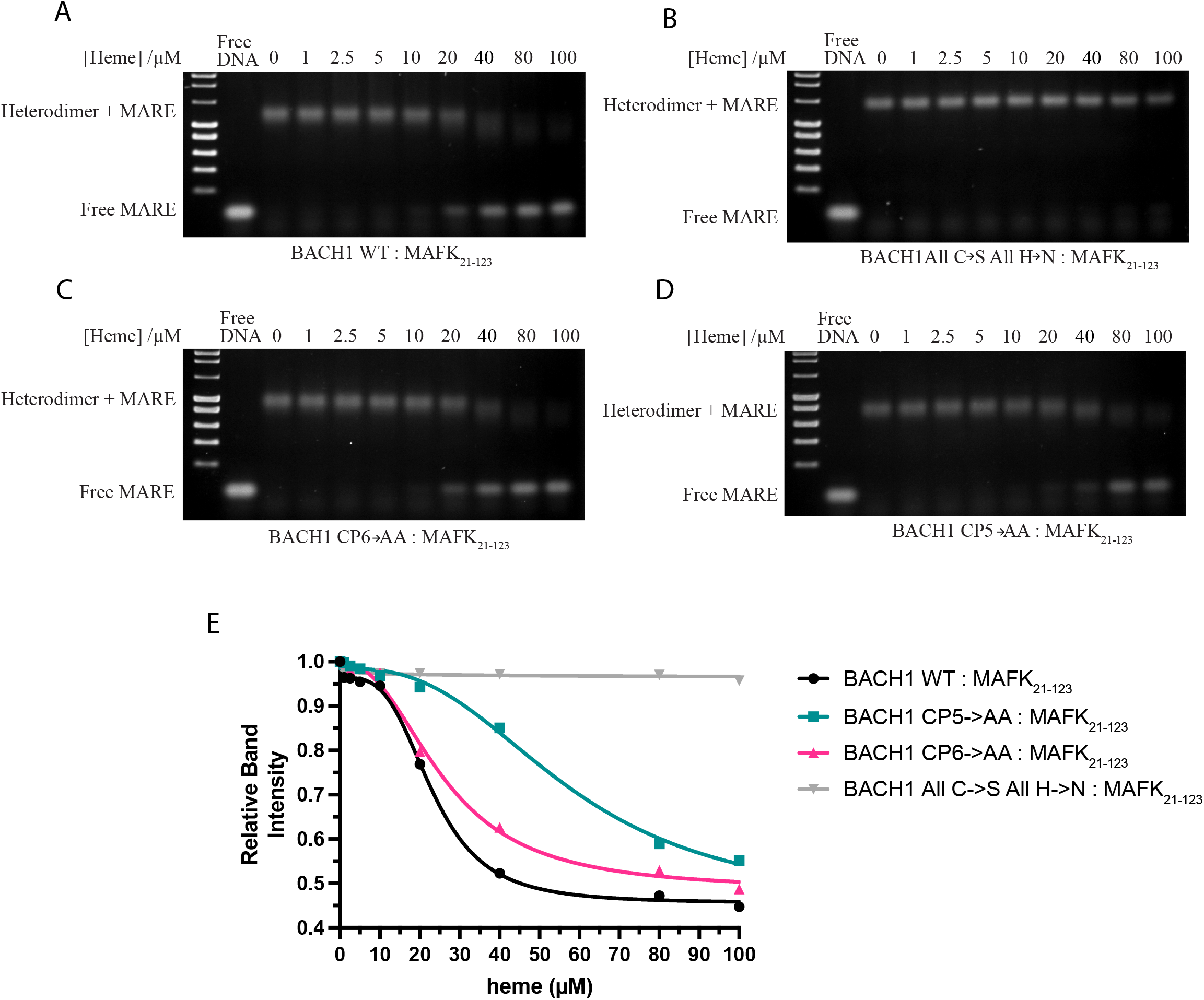
CP5 impaired heme-mediated DNA-dissociation. (A-D) Electrophoretic Mobility Shift assays (EMSA) of WT and different mutant BACH1:MAFK heterodimers (0.5 µM) upon addition of increasing concentrations of heme (0 – 100 µM). Free MARE and heterodimer-MARE complex are indicated. (E) Quantification of free MARE band intensities from (A-D).

Together these results confirmed the expected behaviour of BACH1 and support the involvement of C492(CP5) and H559 in mediating dissociation of the BACH1:MAFK heterodimer from DNA. However, the fact that the C492(CP5) mutation does not abolish heme-mediated dissociation suggests that heme-binding to other (non-CP) ligands is also important.

## Discussion

Our data support the presence of two distinct heme binding pockets in the C-terminal region of BACH1 flanking the DNA-binding domain. NMR experiments suggest that there is a pre-formed heme-binding pocket involving C646(CP6) and C621. UV-visible spectroscopic analysis of BACH1 mutants provide evidence for a second binding site involving C492(CP5) and H559 that is conformationally-induced in the presence of heme. These findings suggest distinct molecular mechanisms for heme-recruitment to the two binding sites in the BACH1 C-terminal region.

### A structured CP6-C621 pocket mediates heme binding but does not contribute to DNA dissociation

Direct evidence for a specific pre-fomred heme-binding pocket involving C646(CP6) and C621 was observed using NMR spectroscopy. Heme addition caused highly specific perturbations in the (^1^H-^15^N)-TROSY spectra of BACH1. Importantly, these changes were lost when either C646(CP6) or C621 was mutated. This indicates that both residues contribute to formation of a high affinity binding site rather than acting as independent coordinate sites for heme. AlphaFold prediction of heme-binding to a BACH1 monomer suggests that the two cysteines could act as axial ligands to the heme iron forming a 6-coordinate heme complex. A binding mode involving two cysteine residues as axial ligands is chemically plausible, as it has been observed in other heme-binding proteins, including the protein DiGeorge Critical Region 8 (DGCR8) (32, 33).

Since BACH1 binds to DNA as a heterodimer with MAFK, it is important to consider whether heme-binding is compatible with a DNA-bound heterodimer. The C646(CP6) and C621 heme-binding pocket is located at one end of the leucine zipper motif, distal to the DNA recognition helix. This means that a BACH1:MAFK heterodimer could bind heme without any disturbance to the leucine zipper heterodimer or the DNA-recognition helix (Figure 3G). This is consistent with our observation that mutation of CP6 in BACH1:MAFK heterodimer has no effect on heme-mediated dissociation from DNA.

Importantly, in the absence of MAFK, BACH1 forms a homodimeric species that modelling suggests could bind two hemes with each monomer engaging C646(CP6) and C621 as axial ligands again resulting in no disturbance to the leucine zipper (Figure 3H). This region has been reported to mediate interaction of a BACH1 homodimer with the CRL2^FEM1B^ E3 ligase, in a heme-dependent fashion, triggering the degradation of BACH1 (26, 34). Since this occurs in the cytoplasm, it makes sense that this involves a BACH1 homodimer rather than the nuclear BACH1:MAFK heterodimer. In contrast to our findings which suggest requirement for both C646(CP6) and C621, Heider et al. modelled a single heme bound between the CP6 ligands from each of the BACH1 monomers (34).

Interestingly, the models of BACH1 suggest that cysteine Cys625 is perfectly positioned to make a covalent bond with one of the heme vinyl groups (Supplementary Figure S7A). This is analogous to heme-bound Cytochrome c (Supplementary Figure S7B) as observed in multiple crystal structures (PDB ID: 2YKZ, 1H21, 3ZOW & 1CNO) (35–38). The interaction of this proximal cysteine with a vinyl group of heme, may both improve the structural stability and prevent heme dissociating from the protein. If this were the case for BACH1, this might effectively commit the protein to degradation. However, we could not detect covalently bound heme by mass-spectrometry of our in vitro purified protein.

### The CP5/H559 site links heme binding to DNA dissociation

UV-visible spectroscopy monitoring of heme binding by various BACH1 mutants suggests a second site can support heme-binding. Constructs containing both C492(CP5) and H559 show strong spectroscopic evidence of a histidine-heme interaction. Constructs lacking either H559 or C492(CP5) lose this interaction suggesting clear cooperation in heme recruitment. The involvement of a histidine in heme-binding is consistent with the findings of Segawa et al. who suggest that the C492(CP5) motif binds heme with a 6-coordinate iron where the sixth ligand is a histidine residue (25). We used AlphaFold predictions to investigate the potential structure around this heme-binding site, which suggests that the C492 (CP5) region may be flexible in the absence of heme. Such flexibility could allow BACH1 to accommodate heme through induced local rearrangements, stabilising an otherwise dynamic region.

DNA binding assays confirmed the expected behaviour of BACH1 and support the involvement of C492(CP5) (likely together with H559) in mediating dissociation of the BACH1:MAFK heterodimer from DNA. However, the fact that the C492(CP5) mutation does not abolish heme-mediated dissociation suggests that heme-binding to other (non-CP) ligands may also be involved.

### BACH1 may employ a broader heme-sensing network rather than a simple CP-motif mechanism

The role of CP sequences as heme-recognition motifs (HRMs) is well established in the heme-regulated transcription factor HAP1, where they are located in intrinsically disordered regions of the protein. In HAP1, heme binding to the HRMs leads to dissociation of chaperone proteins, facilitating DNA-binding and transcriptional activation (21, 22). Our studies suggest that heme-binding to BACH1 works somewhat differently. One heme-binding site is structurally preformed, whereas the other, presumably like those in HAP1 is conformationally-induced on heme-binding. Both binding sites in BACH1 appear to require non-CP ligands to fully coordinate the heme. It is particularly striking that the BACH1 protein contains so many potential heme ligands (34 Cys and 13 His). Of these, 14 cysteines are located in the C-terminal region in proximity to the bZip DNA-binding region. In contrast to BACH1, full-length MAFK contains only one histidine and one cysteine, while full-length NRF2 contains only 6 cysteines. Our UV spectroscopy of the various BACH1 mutant proteins suggests that many, if not all, of these cysteine and histidine residues are competent for heme-binding raising the question as to whether they contribute to the functional recruitment of heme. One possibility is that they act to increase the local concentration of heme facilitating the recruitment of heme to the key functional binding sites.

Interestingly, BACH2, a paralog of BACH1 containing homologous BTB and bZip domains is also cysteine-rich with 38 cysteine and 14 histidine residues. Importantly, the position of the CP motifs and non-CP cysteines in BACH2 is not conserved. Despite this positional drift, the alignment is not simply cysteine loss in one and gain in the other. At several points where BACH1 carries a CP-motif cysteine, BACH2 has cysteine residue at a similar sequence position, even though the proline is lacking. This emphasises that heme-binding may not require a CP dipeptide and approximate, rather than precise positioning may be sufficient to retain function.

Taken together, it is clear that heme-regulation of BACH1 functions through diverse mechanisms and may be more complex than previously thought.

## Materials and Methods

### Cloning and mutagenesis

The BACH1 WT (aa: 477-736), MAFK WT (aa: 21-123) and mutant BACH1 fragments (BACH1 CP5◊AA, BACH1 CP6◊AA and BACH1 CP56◊AA) were cloned into pLEICS expression vector generated by the Protein Expression Laboratory (PROTEX), University of Leicester using a BD In-Fusion^TM^ kit. BACH1 All C◊S All H◊N, BACH1 CP5/6 only, BACH1 other Cys only, BACH1 His only, BACH1 CP6 and C621 only, BACH1 CP5 only, BACH1 H559 only and BACH1 CP5 and H559 only were synthesised and cloned into pLEICS vectors by GeneArt Gene Synthesis Service (Thermo Fisher Scientific). The recombinant proteins were cloned with the following tags, N-GST (for BACH1 WT (aa: 477 to 736)), N-GB1/His6 (for BACH1 WT, BACH1 CP5◊AA, BACH1 CP6◊AA and BACH1 CP56◊AA, MAFK WT (aa: 21-123)), and N-His6/S (for BACH1 All C◊S All H◊N, BACH1 CP5/6 only, BACH1 other Cys only, BACH1 His only, BACH1 CP6 and C621only, BACH1 CP5 only, BACH1 H559 only and BACH1 CP5 and H559 only).

### Protein expression and purification

All constructs were transformed and expressed in *Escherichia coli* Rosetta (DE3) cells at 37°C. The bacterial cells were grown in 2x YT until an OD_600_ of 0.6∼0.8. Protein expression was induced at 20°C for 16-18 hours with 50 µM Isopropyl β-D-1-thiogalactopyranoside (IPTG). Cell pellet was centrifuged at 4 °C at 3,500 x g for 15 min.

N-GST tagged BACH1 WT expressing cells were lysed in 50 mM Tris-HCl pH8, 300 mM NaCl, 5% Glycerol, 5 mM DTT, 1% Triton X-100 and cOmplete EDTA-free protease inhibitor cocktail (Roche). The supernatant fraction was incubated with Glutathionine Sepharose (Cytiva) resin for 1 hour at 4 °C with gentle agitation. The protein-bound resin was washed for 3 times with 50 mM Tris-HCl pH8 buffer containing 300 mM NaCl, 5% Glycerol and 5 mM DTT and then 3 times with 20 mM Tris-HCl pH8 buffer containing 50 mM NaCl, 5% Glycerol and 5 mM DTT. The GST tag was removed by incubating with TEV protease at 4 °C overnight.

All N-GB1-His6 tagged and N-His6-S tagged protein expressing cells were lysed in 20 mM Tris-HCl pH8, 300 mM NaCl, 10 mM Imidazole, 1 mM DTT, 1% Triton X-100 and cOmplete EDTA-free protease inhibitor cocktail (Roche). The supernatant fraction was incubated with Ni-NTA Agarose (Qiagen) for 1 hour at 4 °C. The protein-bound Ni-NTA resin was washed 6 times with wash buffer (20 mM Tris-HCl pH8, 300 mM NaCl, 10 mM Imidazole and 1 mM DTT). All N-GB1-His6 tagged or N-His6-S tagged protein were eluted from the Ni-NTA resin with 300 mM Imidazole. To remove the imidazole and cleave the tag from protein, eluted protein was incubated with TEV protease and dialysed against 4 L dialysis buffer (20 mM Tris-HCl pH 8, 50 mM NaCl and 5 mM DTT) at 4 °C overnight.

All TEV-cleaved proteins were then concentrated using a Amicon® Ultra centrifugal filter concentrator (Merck Millipore) with 10 kDa molecular weight cut off. The concentrated sample was filtered with a 0.22 μm centrifugal filter (Merck Millipore) before loading on a Superdex S200 (10/300) GL column (Cytiva) with Gel filtration buffer (20 mM Tris-HCL pH8, 50 mM NaCl and 5 mM DTT). Fractions were analysed on NuPAGE 4–12% Bis-Tris gels (Invitrogen).

### UV–vis absorption spectroscopy

BACH1 protein samples at a final concentration of 200 µM were incubated with the same molar concentration of heme in Tris-HCl pH 8 buffer, 50 mM NaCl and 5 mM DTT for 10 minutes. Each UV-vis absorption spectrum was scanned using a NanoDrop™ 2000/2000c Spectrophotometer.

### NMR Spectroscopy

For NMR sample preparation, the ^15^N labelled protein sample was incubated with one molar equivalent of heme in 20 mM Tris-HCl pH 8 buffer containing 50 mM NaCl and 5 mM DTT. Preformed heme-bound sample was buffer-exchange into a final NMR buffer (20 mM MES-NaOH pH 6.5, 50 mM NaCl, 5 mM DTT and 5% (v/v) D_2_O). All (^1^H-^15^N)-TROSY spectra were acquired at 298 K using a protein concentration of 250 µM on a Bruker 800MHz or a Bruker AVIII-600MHz spectrometer. Spectra were processed and analysed by using Topspin (Bruker) and Poky® (39).

### Electrophoretic Mobility Shift assays (EMSA)

TRE (TPA responsive element) type MARE (Maf recognition elements) (T-MARE) probe was synthesised and annealed by Integrated DNA Technologies (IDT) using two oligonucleotides, 5’-TCGAGCTCGGAATTGCTGACTCAGCATTACTC-3’ and 3’-TCGAGCCTTAACGACTGAGTCGTAATGAGAGC-5’ (14). To avoid non-specific protein-DNA interactions and non-specific heme-binding, the final buffer composition for each EMSA was 20 mM Tris-HCl pH 8 buffer containing 50 mM NaCl, 5 mM DTT, 0.1 mg/ml BSA, 0.15% Tween-20 and 0.5 µg/µL linear pBR322 vector. To prepare 1:1 heterodimer/DNA complex, 0.5 µM T-MARE was titrated with BACH1:MAFK heterodimer until T-MARE was fully shifted on the agarose gel. The prepared stoichiometric complexes was then incubated with various concentrations of heme (0-100 µM) for 20 mins before loading on a 0.7% agarose, 0.5 x TB gel. The gel was run at 30 mA for 1 hour, and then stained using 5 µg/mL ethidium bromide (EtBr). DNA bands were visualised using UV light. Free MARE band intensities were measured by Image Studio and normalised to 1. Data was plotted against heme concentration. Each curve and pseudo IC50 were generated by using nonlinear fitting in GraphPad Prism.

## Supplementary Figure Legends

**Figure S1: Zoomed-in views of the crystal structures of heme oxygenase (PDB 3I9T) and cytochrome P450 (6IAO).** One DTT and one protein residue form a six-coordinate complex with heme. The distance between the heme iron and side-chain atom (or sulfur atom from DTT) is indicated.

**Figure S2: (^1^H-^15^N)-TROSY spectrum of apo-WT BACH1 (aa: 477-736).** Each resolved cross-peak is numbered from 1-100.

**Figure S3: (^1^H-^15^N)-TROSY spectra of BACH1 (aa: 477-613) and BACH1 (aa: 548-736).** Two truncated BACH1 spectra were used to overlay with the apo-WT BACH1 spectrum (Figure S2) to identify peaks arising from the amino-terminal region and carboxy-terminal region.

**Figure S4: Cross-peak region assignment of apo-WT BACH1 (aa: 477-736).** Cross-peak arising from the amino-terminal region (Green crosses) or carboxy-terminal region (Red crosses) are indicated.

**Figure S5: Overlaid (^1^H-^15^N)-TROSY spectra of apo-WT BACH1 (aa: 477-736) (blue) and mutant BACH1 (CP5/6 --> AA) (red).** Peaks A and B disappeared upon mutation, while peaks C and D appeared in the mutant.

**Figure S6: BACH1:MAFK heterodimer preparation for DNA binding assays.** Size exclusion chromatogram of BACH1:MAFK heterodimer. A260 nm and A280 nm absorbance trace are indicated in red and blue respectively. Protein peaks are indicated by number 1 and 2. **(B)** SDS-PAGE analysis of protein peaks show BACH1:MAFK heterodimer formation. GF load (sample loaded on gel filtration column) and protein peaks as in (A), were loaded on the SDS-PAGE gel. **(C)** WT BACH1:MAFK(aa: 21-123) and mutant BACH1:MAFK (aa:21-123) heterodimer for DNA-binding assays.

**Figure S7: Comparison of heme binding pocket in Cytochrome c crystal structure and AlphaFold 3-predicted models of BACH1. (A)** *Left:* the predicted heme-binding pocket shows C646(CP5) and H559 coordinated to the heme iron, and C625 forms a covalent bond with the heme vinyl group. *Right:* zoomed-in view of heme-binding pocket in CP6 region. **(B)** *Left:* Crystal structure of Cytochrome c with heme (PDB 3ZOW). *Right:* Zoomed-in view of heme-binding pocket showing M61 and H16 coordinated to the heme iron. C12 forms a covalent bond with a vinyl group of heme. The distances between the heme iron and side-chain atom are indicated.

