## Supplementary Figures for "Mechanism of heme binding by CP motifs in the BACH1 DNA-binding region"

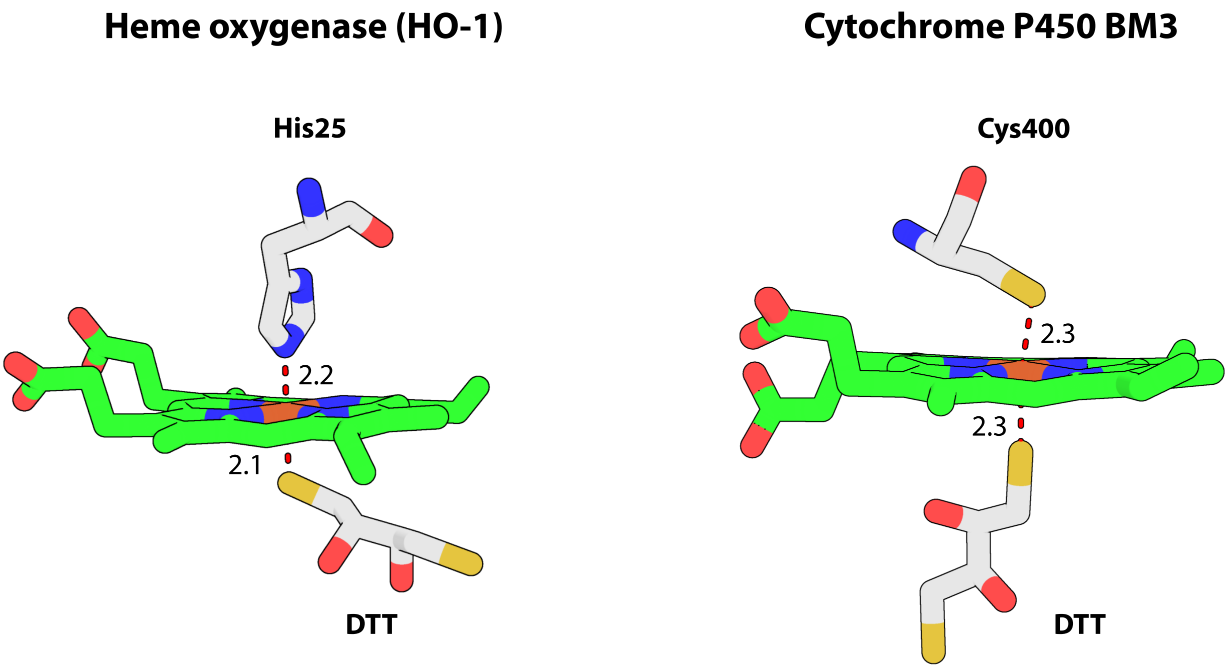


Supplementary Figure S1

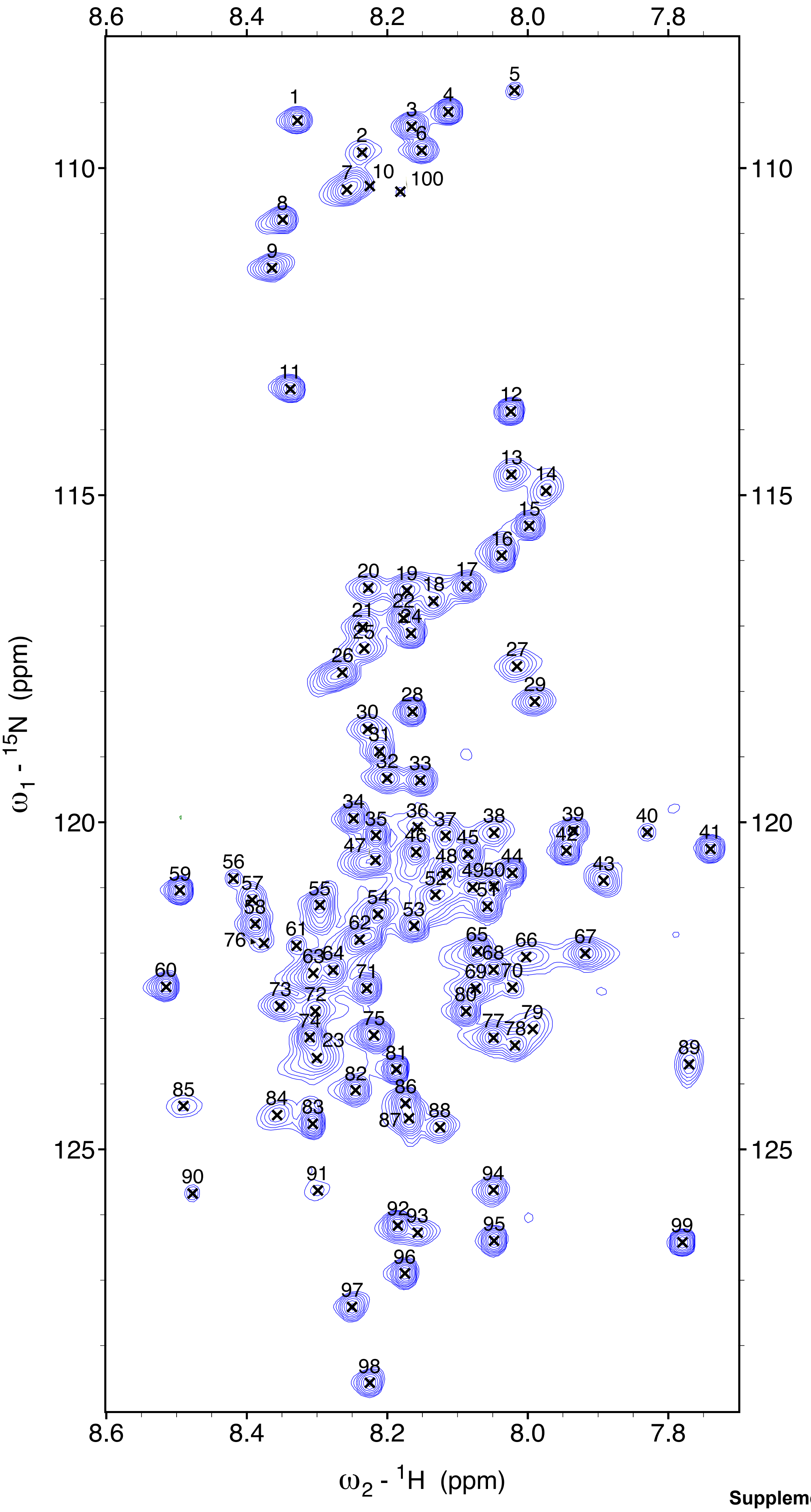


**Supplementary Figure S2**

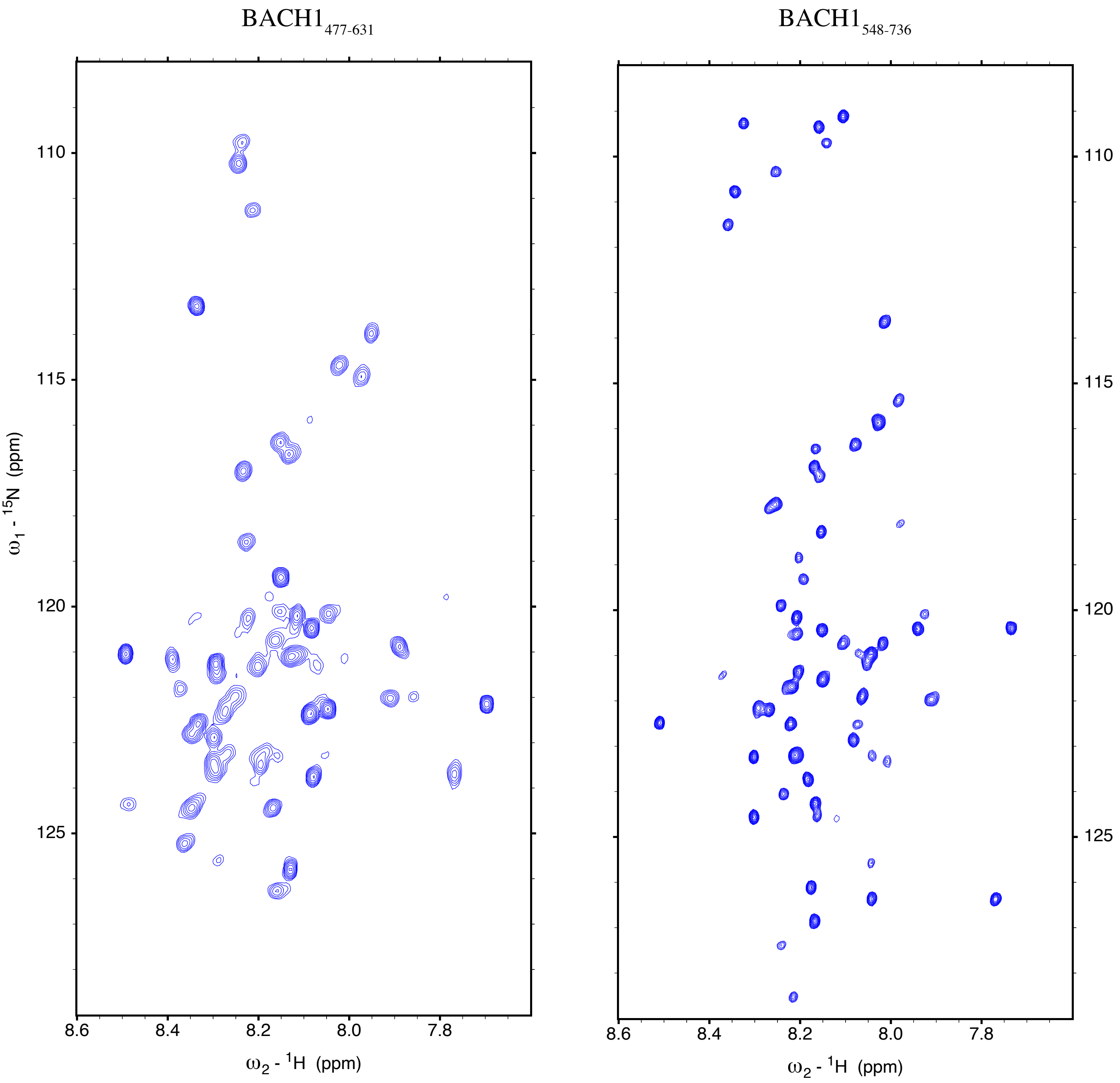


**Supplementary Figure S3**

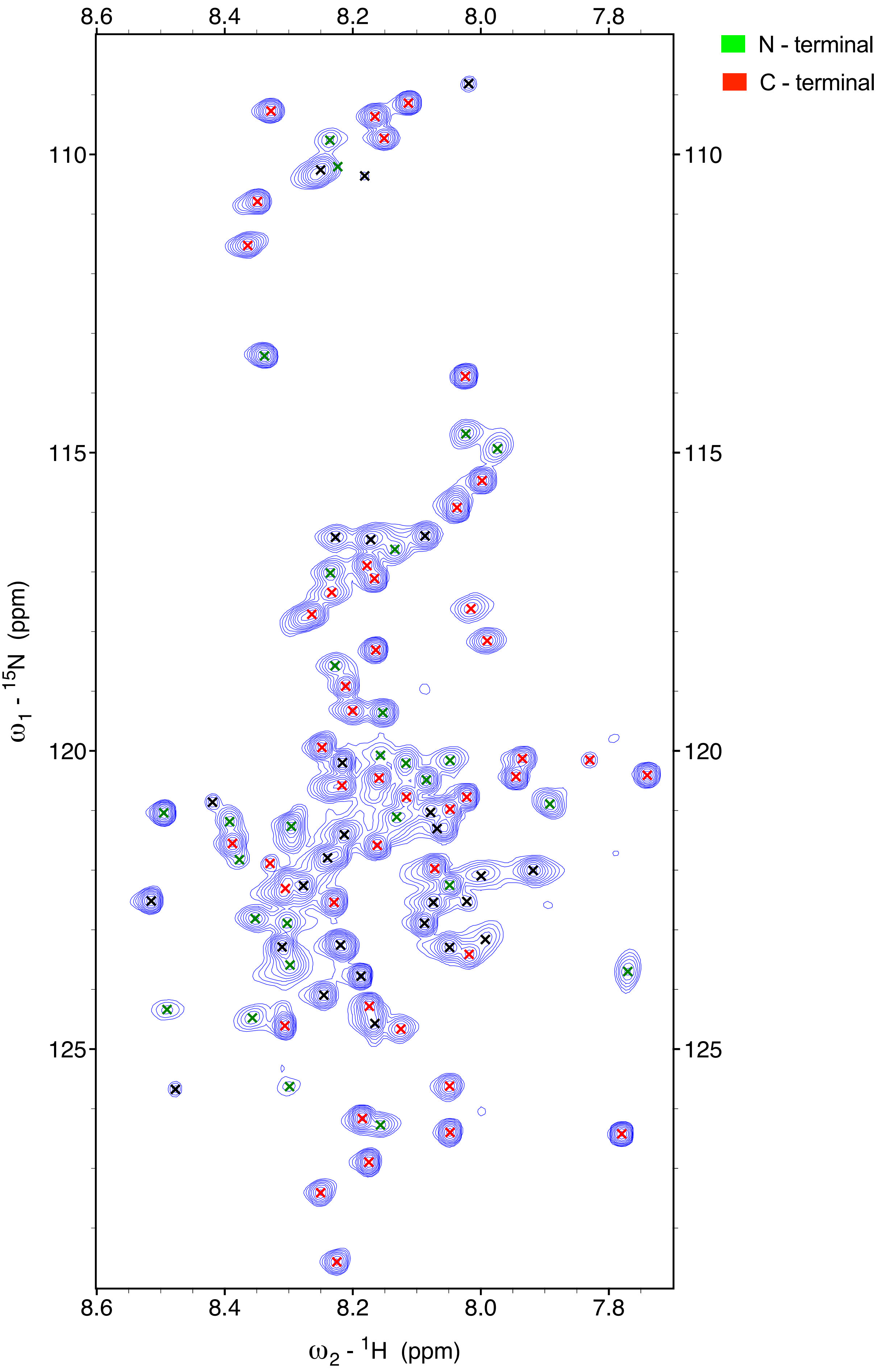


Supplementary Figure S4

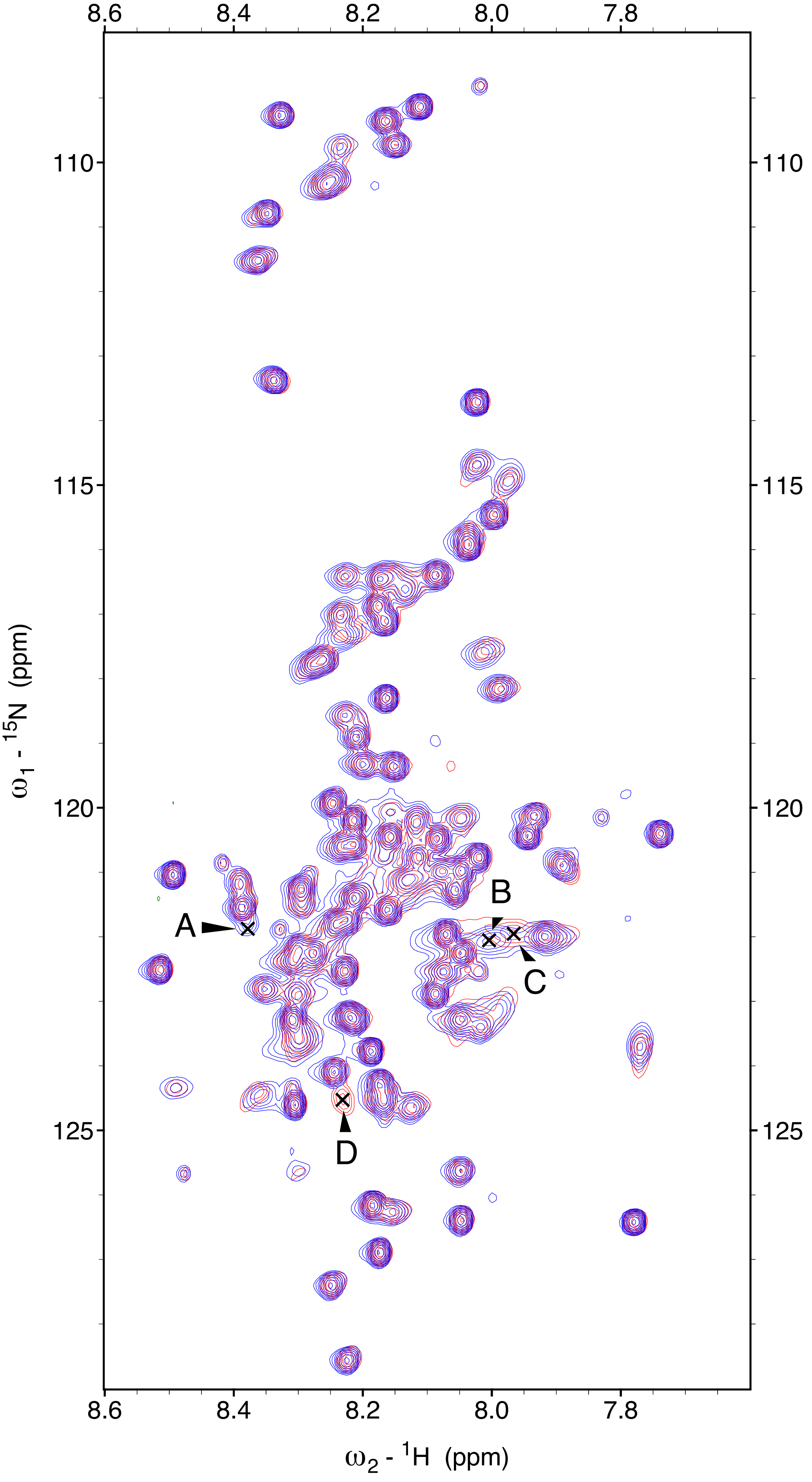


Supplementary Figure S5

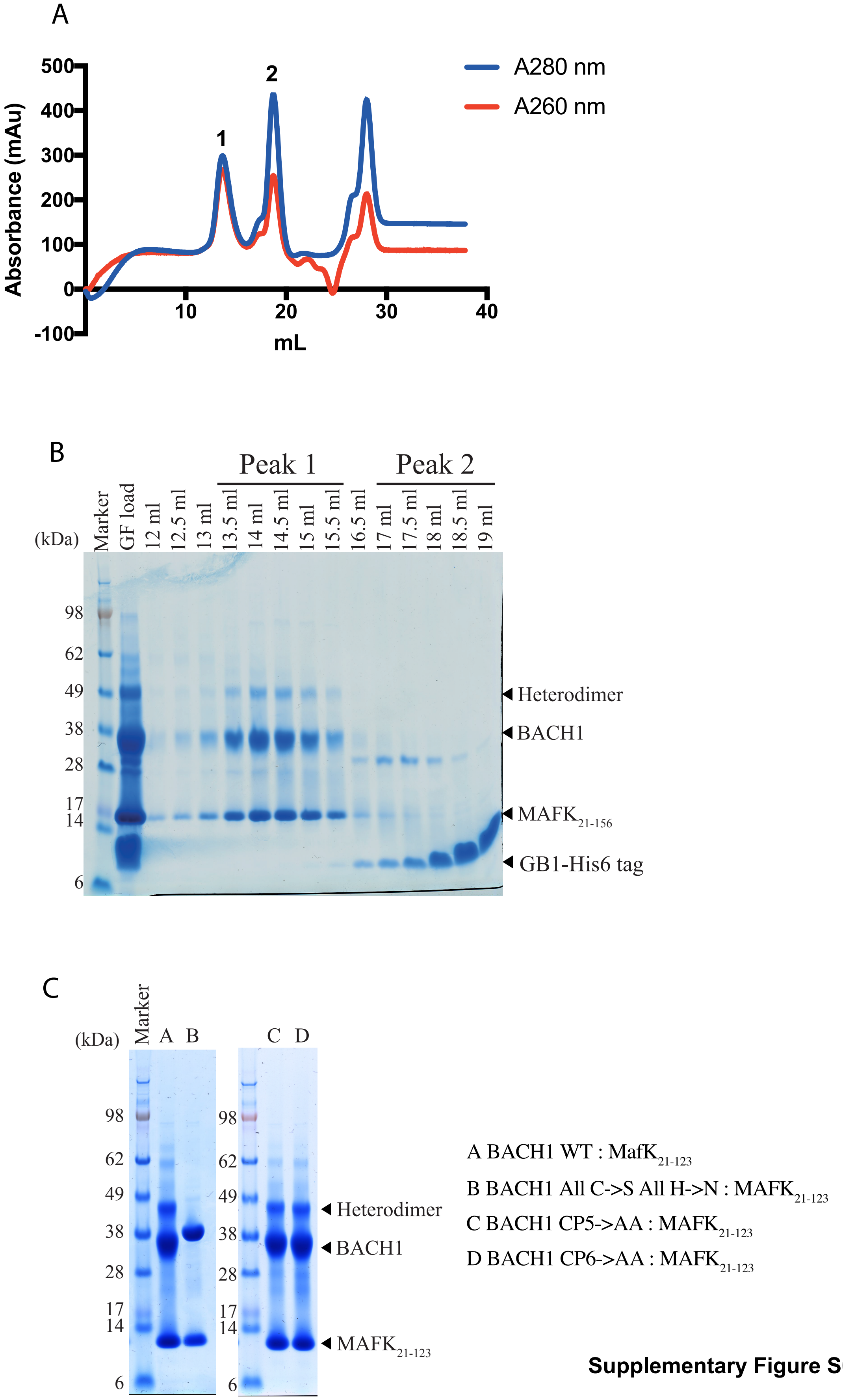


**Supplementary Figure S6**

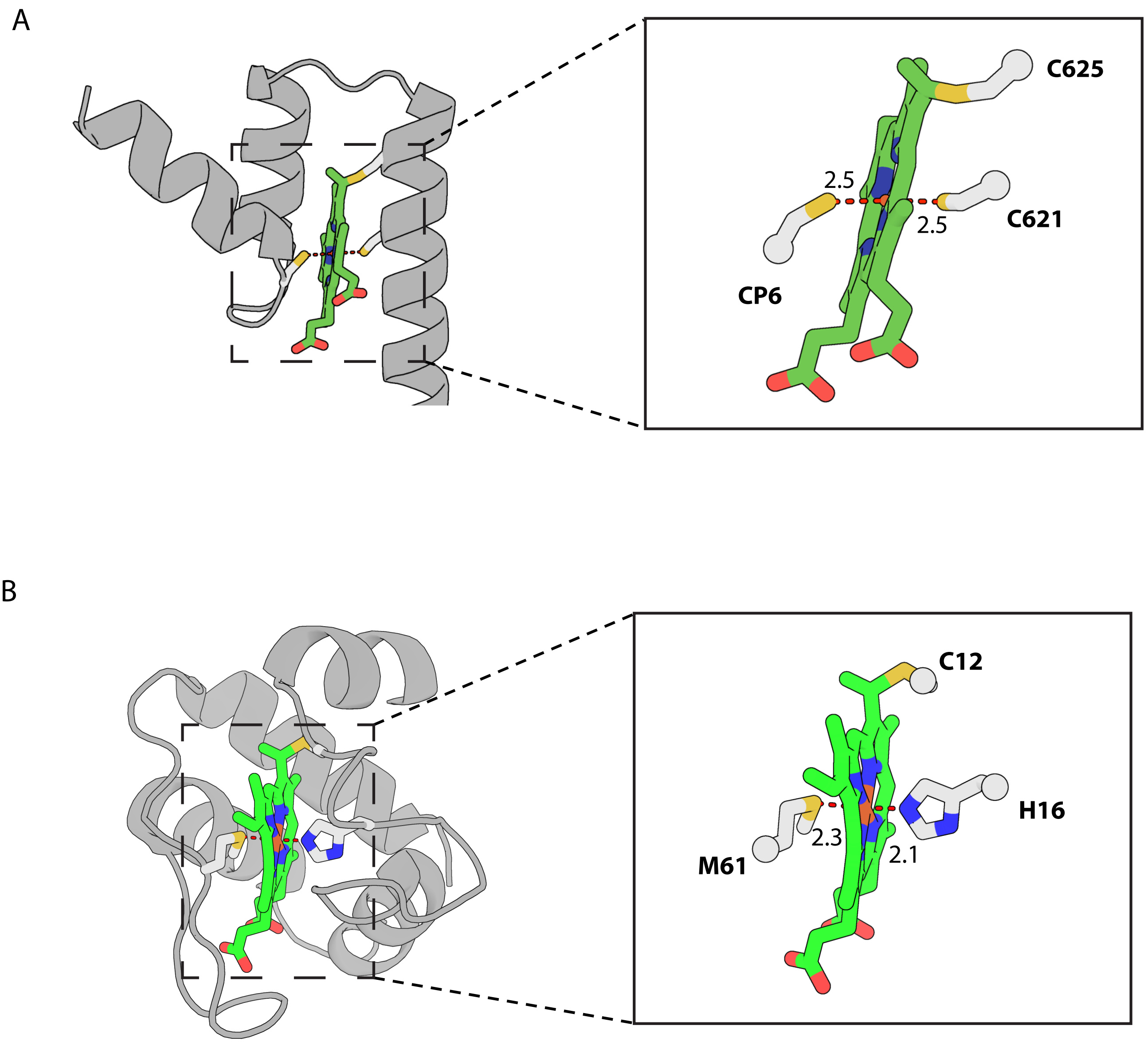


Supplementary Figure S7
